# The Effects of Leukocyte- and Platelet-Rich Plasma (L-Prp) and Pure Platelet-Rich Plasma (P-Prp) an a Rat Endometriosis Model

**DOI:** 10.1101/802421

**Authors:** Ali Doğukan Anğın, İsmet Gün, Önder Sakin, Muzaffer Seyhan Çıkman, Zehra Meltem Pirioğlu, Ahmet Kale, Kayhan Başak, Pınar Kaygın, Serpil Oğuztüzün

## Abstract

Our aim was to investigate the effect of platelet-rich plasma (PRP) derivatives, which can be produced from the patient’s own blood and have minimal side effects, on endometriosis. To the best of our knowledge, this is the first study in the literature that studies the relationship between PRP and endometriosis. Endometriosis foci were created in the first operation. In the second operation (30^th^ day) groups were formed. Group 1 (n= 8) was administered saline, group 2 (n= 7) leukocyte- and platelet-rich plasma (L-PRP), and group 3 (n= 8) pure platelet-rich plasma (P-PRP). Group 4 (n= 10) was used to obtain PRP. In the last operation (60^th^ day), the endometriotic foci were measured, and then excised. There was no statistically significant difference between the pre and post volumes of the endometriotic foci, between their volume differences and volume difference rates (p > .05). However, it was observed that existing implant volumes in all groups decreased statistically significantly within their own groups by the end of the experiment compared to the previous volumes (p < .05). When the implants were assessed through histopathological scoring in terms of edema, vascular congestion, inflammatory cell infiltration, hemorrhage, epithelial line, and hemosiderin accumulation and immunohistochemical staining in terms of VEGF, there was no significant difference in the comparison between the groups. Although L-PRP and P-PRP generated more reduction in the endometriosis foci, they did not create any statistical differences.

## Introduction

Endometriosis, which is described as the presence of endometrial gland and stroma outside the uterine cavity, is an important women’s health problem seen in 6–10% of women that causes degradation in the quality of life with clinical effects, such as infertility, dysmenorrhea, dyspareunia and chronic pelvic pain^1–4^. Its pathophysiology has not yet been fully resolved, and an effective treatment for it has not yet been found^5,6^.

Research has shown that cytokine levels rise in the peritoneal fluid of endometriosis patients^7^. In patients with endometriosis, an angiogenetic activity of peritoneal fluid and increased levels of vascular endothelial growth factor (VEGF) are observed^8,9^. In various experimental studies in the treatment of endometriosis, endometriotic foci have been found to shrink and VEGF levels have been found to decrease^10,11^.

The healing properties of platelet-rich plasma (PRP) and platelet- and leukocyte-rich plasma in tissues have also been subject to numerous research studies in recent years. This plasma contains a high proportion of platelets. Platelets are also known to contain many growth factors. Platelet-derived growth factor (PBGF), transforming growth factor beta (TGF-B), epidermal growth factor (EGF), insulin-like growth factor (IGF) and vascular endothelial growth factor (VEGF) can be counted among these factors^12,13^. With such features, PRP can show positive effects on many systems. Such effects of it include many systems such as scalp, skin, heart, bones, cartilage, tendons, liver, kidney, genital tract, ovaries, endometrium and infertility treatments^14–20^. PRP can be in two different forms: L-PRP (i.e., leukocyte- and platelet-rich) and P-PRP (leukocyte-poor or pure platelet-rich). Although they are similar products, their contents such as cytokines and growth factors are different. L-PRP has a higher proportion of leukocyte, TNF-a and IL-1β concentration^21^. To the best of our knowledge, there is no study in the literature investigating whether PRP administration increases or decreases endometriosis.

Our aim was to investigate the effect of two forms of PRP (L-PRP and P-PRP) on endometriosis, which had never been administered in endometriosis, but was known to be effective in many areas.

## Materials and Methods

The study was carried out in the Animal Experiments Laboratory, and approval was received from UNIVERSITY OF HEALTH SCIENCES Hamidiye Animal Experiments Local Ethics Committee (No:46418926-605.02 Date: 2018-01/01, 2019-01/19)

For the experiment, 34 4-month-old, 250-300 gr, Sprague Dawley type female rats were used.

### First operation: Creation of an endometriosis model

The rats (n= 24) were administered 10% ketamine (80 mg/kg Ketalar; Eczacibasi, Istanbul, Turkey) and 2% xylazine chloride (15 mg/kg, Rompun; Bayer Health Care LCC, Kansas, KS) intraperitoneally for anesthesia prior to laparotomy. Abdomens were shaved and cleaned with iodine (Povidone-iodine 10% solution, Batticon; Adeka Laboratories, Istanbul, Turkey), and each abdomen was entered through a 5-cm vertical incision. As defined by Vernon and Wilson, foci were formed by implanting the part taken from the rat uterus to the abdominal wall through a surgical intervention using Uygur’s modification^22,23^. To do that, a .5 cm section of the left uterine horn was excised at a distance of 1 cm from the ovary. The remaining uterine horn was sutured with 2/0 polyglactin absorbable suture, and the bleeding was controlled. The tissue that was taken was cut longitudinally and sutured without separating the myometrium into the right abdominal peritoneal inner surface with 5/0 polypropylene non-absorbable suture by placing the endometrial portion inward, and an implantation was achieved (to ensure the endometriosis model) (Fig 1). The implants were washed with 5 cc saline flush to prevent possible adhesions and dryness. The abdomen was closed by suturing the peritoneum, fascia and skin with 4/0 polyglactin. After the operation, 50 mg/ kg/ day Cefazolin sodium (IE Ulagay Ilac Sanayi, Istanbul, Turkey) was administered intraperitoneally for 3 days. Each rat was operated in 20 minutes in order to prevent the room air temperature from disturbing the dryness of the tissue. The rats were caged individually in a controlled environment (at 21 °C room temperature and 60% humidity) with 12 h light/dark cycles, and were fed ad libitum.

**Fig 1:**
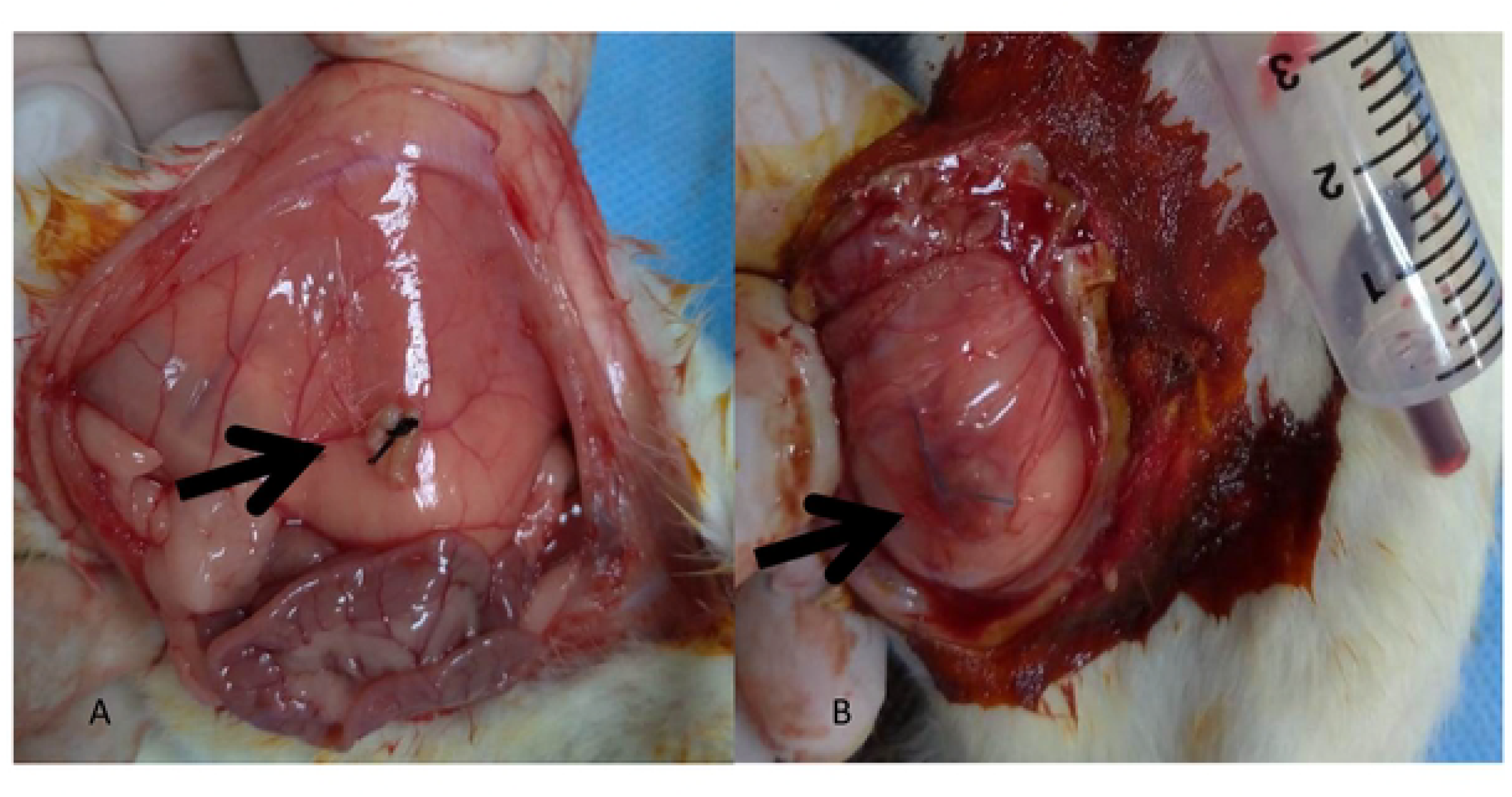
The endometrial focus on the inner abdominal surface of the rat A: Endometrial focus implantation, 1^th^ day. B: Endometriosis implant, 30^th^ day.

### Second operation: Creation of the groups

The second laparotomy was administered 1 month later in order to assess the presence of endometrial foci, their transformation into a cystic structure, and their dimensions. The abdomen was entered through the previous incision (anesthesia, cleaning and antibiotics were administered in the same way as in the first operation). The implants were found to be successful in all rats (Fig 1). The implant dimensions were measured and the global endometriotic focal volumes of the implants were calculated using the prolate ellipsoid formula (V mm^3^= .52× A× B× C where A, B, and C are width, length, and height, respectively)^24^.

The rats were divided into 3 groups with random selection:

Group 1: Control group (n= 8): 0,1 cc SF was applied on the implant.
Group 2: Leukocyte- and platelet-rich L-PRP group (n= 7): 0,1 cc L-PRP was applied on the implant.
Group 3: Pure platelet-rich P-PRP group (n= 8): 0,1 cc P-PRP was applied on the implant.

A total of 10 rats were decapitated after the blood was drawn for the preparation of the heterologous PRP. All the injections were applied once on the lesion in all groups. After that, the abdomen was closed by suturing the peritoneum, fascia and skin with 4/0 polyglactin. A rat in Group 2 died 3 days later, and there were 7 rats remaining in the group.

### Third operation: Termination and pathological examination

A laparotomy was performed for the third and last time, for final assessments 1 month later. In the last 5 days, vaginal smears were sampled from all rats to assess estrogen cycle. The cycle status was assessed by microscopic examination through the Papanicolaou staining method. The vaginal smears were taken in the form of swabs from the vaginal wall by using a cotton brush. The estrogen cycle was determined by the cornification of the cells formed by the estrogen effect and loss of leukocytes^24^. The rats that were in their cycles were selected. The pre-operative anesthesia and cleaning were performed as before. The abdomen was entered through the previous incision line. The endometriosis foci were measured by the same researcher using the same method (the prolate ellipsoid formula) as stated above, blindly by not knowing which group the foci were in. After that they were excised. Then, the rats were decapitated (cardiac excision) and were destroyed by red medical waste bins. The tissues that were excised were sent to the laboratory within 10% formaldehyde for histopathological and immunohistochemical examination. The pre- and post-treatment implant volumes, post-treatment histopathological examination scores of the implants and immunohistochemical staining scores for the post-treatment VEGF in the implants were measured and compared.

### Preparation of PRPs

Ten additional rats were used to obtain blood for PRP. These rats (n= 24) were administered 10% ketamine (80 mg/ kg Ketalar; Eczacibasi, Istanbul, Turkey) and 2% xylazine chloride (15 mg/ kg, Rompun; Bayer Health Care LCC, Kansas, KS) intraperitoneally for anesthesia, and their blood samples were drawn through cardiac puncture. The blood was anticoagulated using acid-citrate dextrose solution A (ACD-a) at a rate of 1/9. A total of 38-40 cc PRP (L-PRP and P-PRP) was obtained from the 10 rats.

### Preparation of L-PRP

L-PRP was prepared using the double centrifuge method based on buffy coat. The whole blood from five rats was centrifuged at room temperature for 10 minutes at 250 g, and it was ensured that the blood was separated into three layers: Erythrocytes at the bottom, buffy coat in the middle (rich in platelets, leukocytes and fibrinogen), and plasma containing platelets at the top. The platelets-containing plasma and buffy coat were later transferred into a new tube. A large portion of the platelets, leukocytes, and fibrinogen was re-centrifuged for 10 minutes at 1000 g to form precipitate. The supernatant plasma was thrown away, and the precipitated platelets were re-suspended in the residue plasma to obtain L-PRP^25,26^.

### Preparation of P-PRP

P-PRP is a plasma-based method that concentrates platelets and eliminates leukocytes and erythrocytes. The anticoagulated whole blood that was drawn from the five rats was centrifuged at room temperature for 10 minutes at 160 g to separate platelets-containing plasma (rich in leukocytes) from erythrocytes and the buffy coat. Attention was paid to prevent the buffy coat and erythrocyte contamination. The platelets-containing plasma was then transferred to a new tube and centrifuged for 15 minutes at 250 g. The supernatant plasma was thrown away, and the precipitated platelets were re-suspended in the residue plasma to obtain L-PRP^25,26^.

### Histopathological examination

All pathological examinations were blindly carried out by a single expert (K.A.). Biopsies were fixated for 24 hours in 10% formaldehyde. Paraffin blocks were created, and the blocks were cut in thickness of 5 um. A total of 5 sections were obtained for each material, stained with hematoxylin eosin (HE) and assessed with a light microscope. Edema, vascular congestion, inflammatory cell infiltration, fresh hemorrhage and hemosiderin formations were noted (scoring 0–3 where 0= None, 1= Light, 2= Medium, 3= Heavy). Histopathological diagnosis was determined by the recognition of endometrial tissue, gland and stroma, and by the determination of endometrial lining and luminal formation. The presence of endometrial cells in autografts was assessed semi-quantitatively. The pathological evaluation of the uterine autografts was carried out as described in an earlier method — A well-preserved epithelial line= 3 points, a moderately preserved epithelium with leukocyte infiltration= 2 points, a poor epithelium with rare epithelial cells= 1 point, and no epithelium= 0 points^24^ (Fig 2).

**Fig 2:**
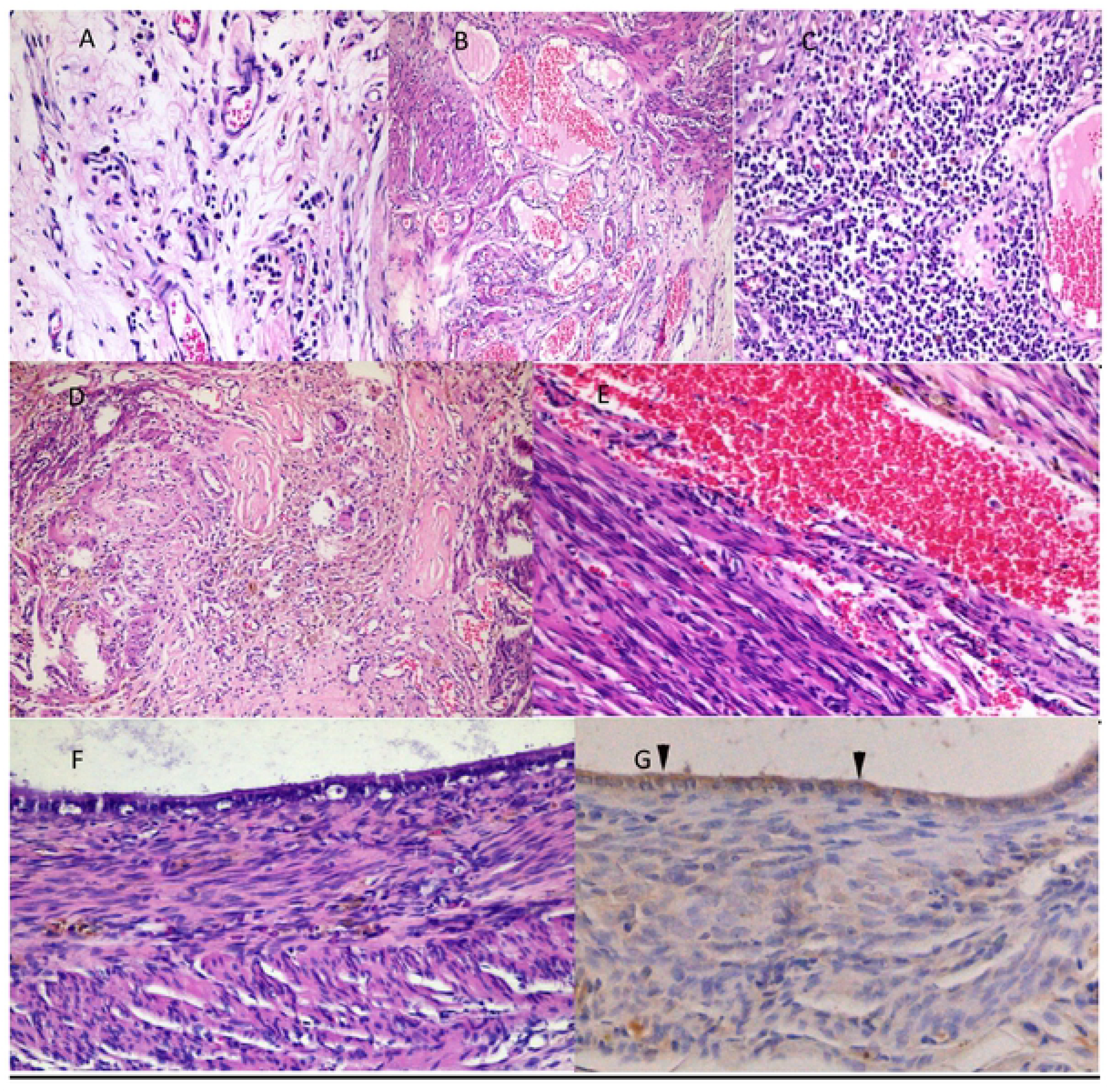
Histopathological appearance and immunohistochemical staining of endometrial implants A: positive edema (H&E × 400) B: Numerous dialted vascular structures (H&E × 200) C: Dense mixed inflammatory cells (H&E × 400) D: Diffuse hemosiderin pigment (H&E × 200) E: Extensive new bleeding area (H&E × 400) F: VEGF positivity in epithelial line (× 400) G: Epithelial line (H&E × 400)

### Immunohistochemical Staining

Tissues were fixed in 10% buffered formalin and embedded in paraffin blocks. Sections that were 4 μm thick were cut, and one section was stained with haematoxylin-eosin to observe the tissue morphology. For immunohistochemistry, endogenous peroxidase activity was blocked by incubating the sections in 1% hydrogen peroxide (v/v) in methanol for 10 minutes at room temperature (RT). The sections were subsequently washed in distilled water for 5 minutes, and antigen retrieval was performed for 3 minutes using 0.01 M citrate buffer (pH 6.0) in a domestic pressure cooker. After washing in distilled water, the sections were transferred in 0.05M Tris-HCl (pH 7.6) containing 0.15 M sodium chloride (TBS). The sections were incubated at RT for 10 minutes with super block (SHP125) (ScyTek Laboratories, USA) to block nonspecific background staining. The sections were then covered with the primary antibodies diluted 1:25 for anti-VEGF in TBS at 4°C overnight (Anti-VEGF (Novus Biologicals NB100-698) After washing in TBS for 15 minutes, the sections were incubated at RT for biotinylated link antibody (SHP125) (ScyTek Laboratories, USA). Then, treatment was followed with Streptavidin/HRP complex (SHP125) (ScyTek Laboratories, USA). Diaminobenzidine was used to visualise peroxidase activity in the tissues. Nuclei were lightly counterstained with haemotoxyline, and then the sections were dehydrated and mounted. Both positive and negative controls were included in each run.

Immunoreactive cells were recorded during the immunohistochemical examination for VEGF with the following scoring: 0=negative staining, 1= < 33% positive staining, 2= 33– 66% positive staining, 3= > 66% positive staining (Fig 2). The immunohistochemical staining was evaluated by the same histologist blindly by a semi-quantitative method using the H-score. For each section, positive areas were scored at × 400 magnification from 0 to 3+ with no staining (0), weak (1+), moderate (2+), and strong (3+). H-score was calculated as H= Ʃ Pi (I + 1). ‘Pi’ represents the density of immunohistochemical staining and ‘I’ is the percentage of the stained cells^10^.

## Results

Three groups were formed in the study — Group 1: Control, Group 2: L-PRP, and Group 3: P-PRP. It was confirmed by the pathologist that the foci were histopathologically endometriosis in all groups. Among the groups, the pre and post volumes of the endometriotic foci created, volume differences between them and volume difference rates between them are seen in Table 1. Considering this table, there is no statistically significant difference between the groups (p > .05). However, it was observed that existing implant volumes in all groups decreased statistically significantly within their own groups by the end of the experiment compared to the previous volumes (p < .05) (Fig 3). When the implants were assessed through histopathological scoring in terms of edema, vascular congestion, inflammatory cell infiltration, and fresh hemorrhage, there was no significant difference in the comparison between the groups in terms of the total score that was obtained (Table 1).

**Table 1:** Comparison of histopathological total score and volume values

| parameters | Group 1<br>(n=8) | Group 2<br>(n=7) | Group 3<br>(n=8) | TOTAL | p |
| --- | --- | --- | --- | --- | --- |
| First volume | 61.95±54.76 | 37.67±26.04 | 29.00±13.60 | 43.10±37.53 | .616<sup>c</sup> |
| **Last volume** | 11.05±15.37 | 2.38±4.50 | 6.79±9.56 | 6.93±11.07 | .228<sup>c</sup> |
| **Volume difference** | 50.90±54.53 | 35.29±27.93 | 22.20±14.16 | 36.16±37.05 | .314<sup>a</sup> |
| **Percentage of difference** | 67.86±68.11 | 88.40±22.03 | 79.41±30.61 | 78.13±44.49 | .384<sup>c</sup> |
| **Total score\*** | 3.88±1.36 | 4.86±1.57 | 3.63±1.51 | 4.09±1.50 | .264<sup>a</sup> |
| **p** | **.025<sup>b</sup>** | **.018<sup>b</sup>** | **.012<sup>b</sup>** | **< .001<sup>b</sup>** |  |

The data are given as average ± standard deviation. \*Total score: Edema + vascular congestion + inflammatory cell infiltration + fresh hemorrhage. <sup>a</sup>ANOVA, <sup>b</sup>Wilcoxon Signed Ranks, first volume and last volume comparison, <sup>c</sup>Kruskal-Wallis

**Fig 3:**
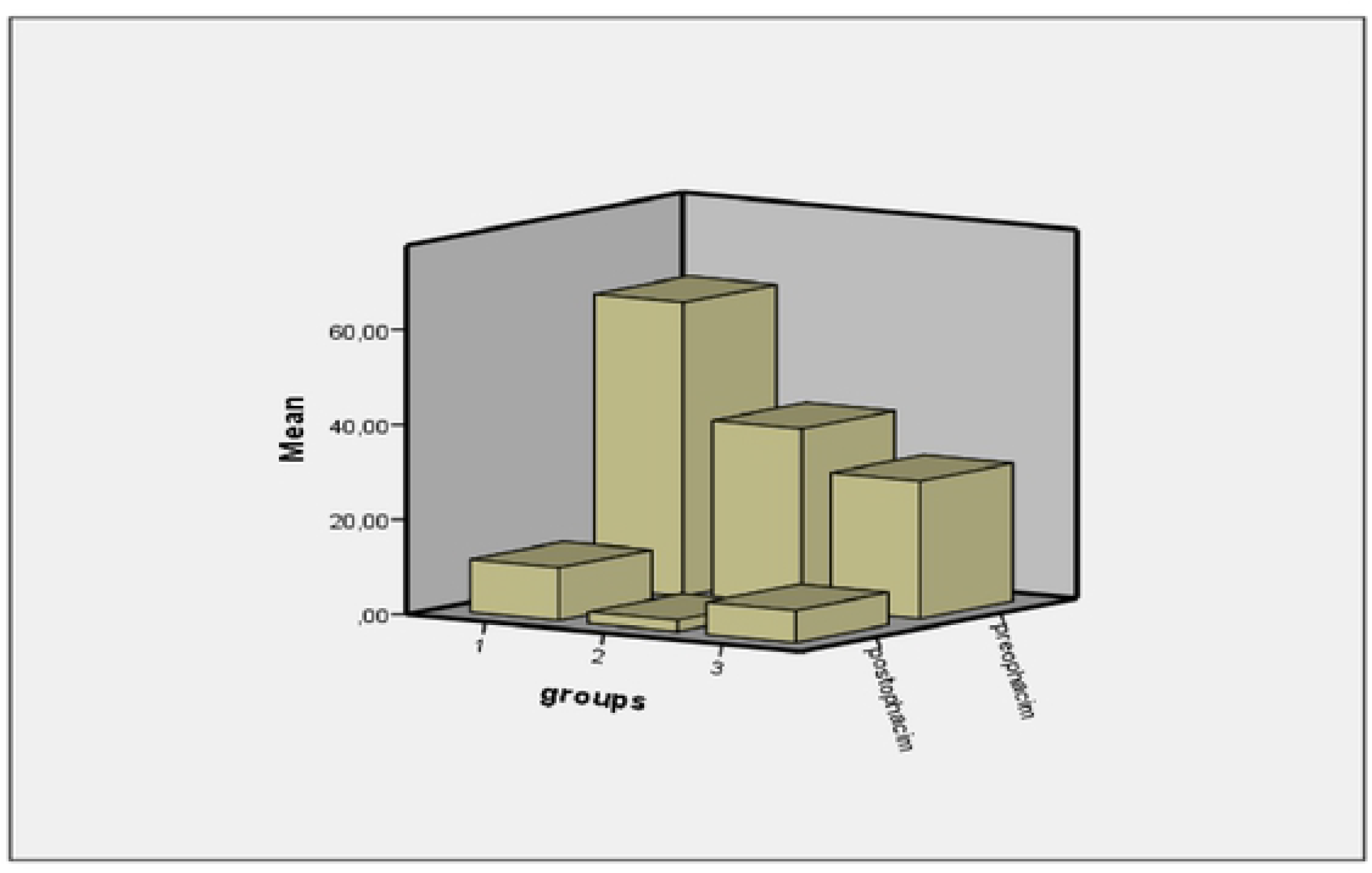
Pre and post implant volumes

No significant differences were found when the groups were compared in terms of the percentages of VEGF score measured immunohistochemically, the percentages of epithelial line score used to evaluate the presence of endometriosis, and the percentages of score indicating hemosiderin accumulated in the implants (Table 2) (p > .05).

**Table 2:**
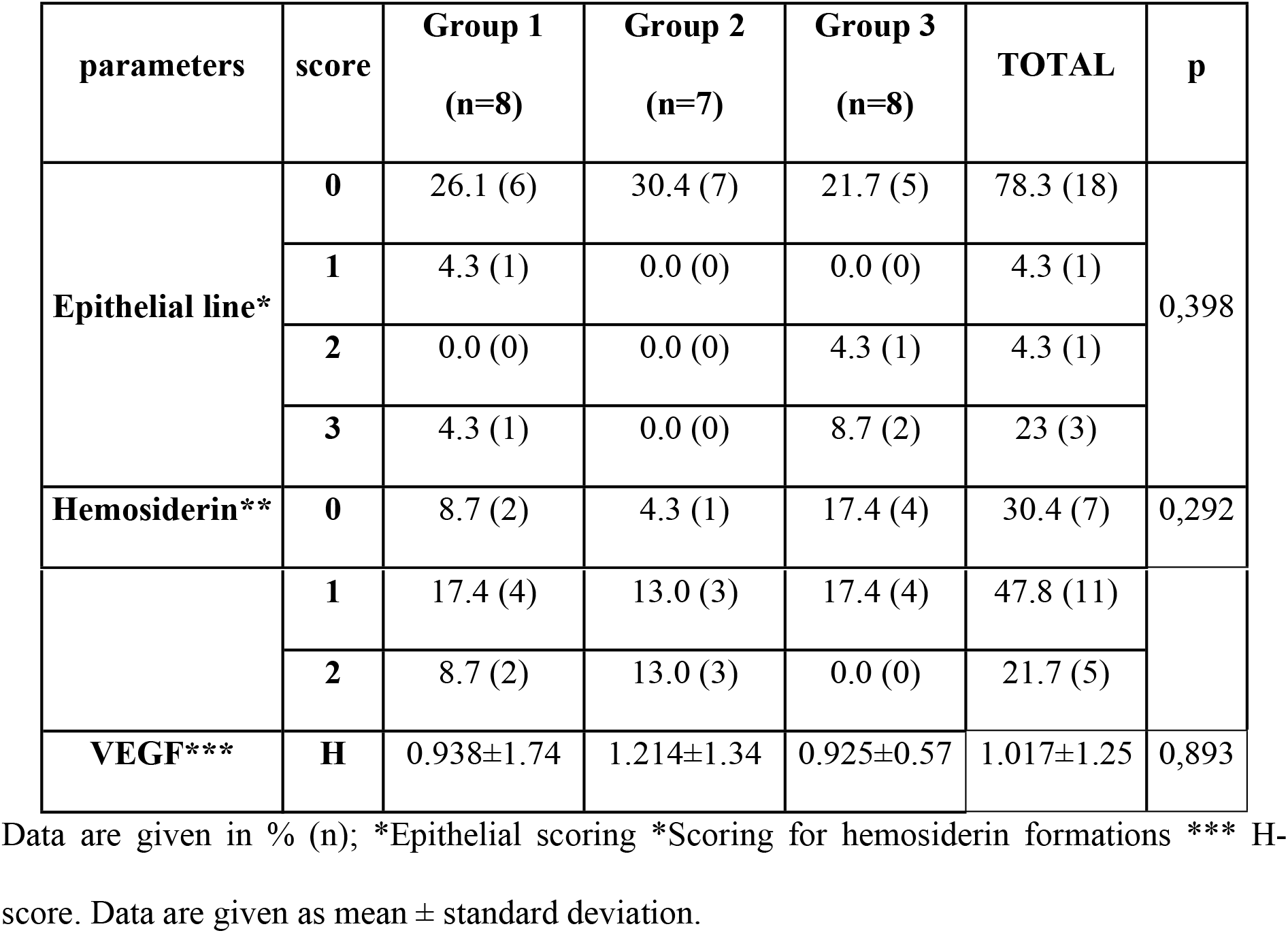
Comparison of histopathological and immunohistochemical parameters

| **parameters** | **score** | **Group 1**<br>**(n=8)** | **Group 2**<br>**(n=7)** | **Group 3**<br>**(n=8)** | **TOTAL** | **p** |
| --- | --- | --- | --- | --- | --- | --- |
| **Epithelial line\*** | **0** | 26.1 (6) | 30.4 (7) | 21.7 (5) | 78.3 (18) | 0,398 |
|  | **1** | 4.3 (1) | 0.0 (0) | 0.0 (0) | 4.3 (1) |  |
|  | **2** | 0.0 (0) | 0.0 (0) | 4.3 (1) | 4.3 (1) |  |
|  | **3** | 4.3 (1) | 0.0 (0) | 8.7 (2) | 23 (3) |  |
| **Hemosiderin\*\*** | **0** | 8.7 (2) | 4.3 (1) | 17.4 (4) | 30.4 (7) | 0,292 |
|  | **1** | 17.4 (4) | 13.0 (3) | 17.4 (4) | 47.8 (11) |  |
|  | **2** | 8.7 (2) | 13.0 (3) | 0.0 (0) | 21.7 (5) |  |
| **VEGF\*\*\*** | **H** | 0.938±1.74 | 1.214±1.34 | 0.925±0.57 | 1.017±1.25 | 0,893 |

Data are given in % (n); \*Epithelial scoring \*Scoring for hemosiderin formations \*\*\* H-score. Data are given as mean ± standard deviation.

### Statistical Analysis

For statistical studies, the IBM SPSS statistics version 24 was used. The Shapiro-Wilk and Kolmogorov Smirnov tests were used to test the normality of distributions. The one-way analysis of variance (ANOVA) was used when comparing three or more groups with a normal distribution, whereas the Kruskal Walls test was used when comparing three or more groups with a non-normal distribution. Following that, the Mann Whitney U test with Bonferroni correction was used for pairwise comparisons. The Chi-square test was used when comparing categorical variables. Paired samples t-tests or the Wilcoxon test were used depending on the conditions in repeated pairwise measurements at different times in the dependent groups. A p value less than .05 was considered as statistically significant.

## Discussion

Endometriosis, which has an important place in female infertility, and whose treatment and pathophysiology are still not certain, has been considered a serious disease today. With the surgery applied in endometriomas, infertile patients face a risk of reduction of ovary reserve^27,28^. Surgical interventions in particularly deep endometriosis can lead to serious complications^29^. Therefore, noninvasive therapies are noteworthy. In this case, PRP, which is safe because it is produced from the patient’s own blood^30,31^ and may be an alternative to surgery and other medications with many side effects, as a minimally invasive agent in the treatment of endometriosis.

There are many studies on endometriosis models and the effects of different drugs in rats. In general, comparisons in these studies have been performed based on volumes prior to and after drug administration, histopathological scores, and various immunohistochemical and biochemical assessments. In the rat experiment, where Yıldırım et al. examined the effects of etanercept on endometriosis, they detected significant reduction in the pre- and post-treatment focal volumes in the pharmaceutical group^32^. No reduction was observed in the control group. They observed that the volume of the endometriotic foci had shrunk spontaneously within 6 weeks in the 2nd control group which did not receive any medication. However, they did not evaluate the rate of volume change between the groups. Moreover, they administered estrogen in certain periods in all groups except for the 2nd control group^32^. Islimiye et al. also carried out a similar experiment with etanercept, but they did not use estrogen; they found that the volume of the implant increased in the control group, decreased in the etanercept group and that this change was significant compared to the control group^24^. In another study, again, where estrogen was not used, the volume after the treatment was significantly less compared to the control group in resveratrol and leuprolide acetate groups^10^. We also found in our study, where we did not administer estrogen, that implant volumes in PRP groups were significantly decreased after the treatment (p < .05). However, this significant decline was similarly present in the control group, and the rate of volume change did not show any significant difference between the groups (Table 1).

Another parameter that is compared in endometriosis studies is the histopathological evaluation of the endometrial glandular and stromal structures that are carried out semi-quantitatively. In a number of studies, there have been significantly lower values compared to the control group after various treatments, while in some others, there have been no significant changes^10,24,32^. We did not observe any significant differences in the post-treatment groups in which we carried out epithelial assessments similar to the studies in the literature (Table 2). However, the point is that in the L-PRP group, the epithelial score was 0. There were no cells. L-PRP had almost destroyed the endometrial foci. Nevertheless, this circumstance had not been reflected in the accumulation of hemosiderin. It was similar in every group (Table 2). Hemosiderin is a significant indicator in the assessment of endometriosis^33,34^. In other words, it seems that the endometrial focus examination should not be carried out based on a single factor. We also assessed the inflammatory changes in histopathological scoring as a total score in our study but did not observe any differences between the groups (Table 1).

While there are many studies of PRP, which includes several growth factors, in different disciplines, studies conducted in the field of gynecology and obstetrics are limited in the literature. It has been stated that PRP contributes to the endometrial growth and thickening and may be effective in infertility in patients with a thin and weak endometrium^35–37^. Farimani et al. have stated that local PRP administration prior to embryo transfer in recurrent implantation failures may improve the success of implantation^38^. PRP can also suppress the inflammatory process in the development of endometriosis^39^.

PRP with its mitogenic effect is important in the renewal and repair of tissues; it does this through its growth factors such as dense platelet-related TGF-B (transforming growth factor-β) and VEGF, and cytokines^40^. In conclusion, PRP seems to be suppressing the inflammatory process^41^. TGF-B is one of the cytokines involved in adhesion pathophysiology^42^. In the study of Murat et al., adhesions were decreased after the PRP administration; additionally, the TGF-B expression in the adhesion foci where PRP was administered has been shown to decrease^25^. This means that although PRP contains TGF-B, it both reduced adhesion and decreased TGF-B expression in adhesions. In another study on the healing of femoral avascular necrosis, TGF increased significantly in the PRP group^43^. VEGF is a cytokine that has a role in angiogenesis and is involved in the pathophysiology of endometriosis^44^. Resveratrol and similar drugs that inhibit the release of VEGF have been shown to reduce endometriosis and cause decreased levels of VEGF in foci^45,46^. Although it has been shown that VEGF levels increase in the tissue with PRP treatment, there are also studies that show that there is no increase and that the treatment does not cause any difference^47–49^. We did not see any significant difference between the VEGF levels in the post-treatment implants in our study, either (Table 2). The tissues in which the effects of PRP have been examined in the literature are different tissues of the body, and perhaps the reason for this difference in the studies may be due to the possibility that the effect of PRP varies depending on the tissue. Therefore, other studies to be carried out in similar tissues are needed.

There are also cytokines such as IL-1, IL-8 in PRP^50^. Marini et al. showed that PRP reduced IL-1B and IL-8 release in endometrial tissues and they attributed the anti-inflammatory effect of PRP to this reduction^51^. Some of the cytokines held responsible for the pathophysiology, which are shown to increase in the peritoneal fluid in endometriosis, are IL-1 and IL-8^44^. That is, although PRP contains IL-1 and IL-8, it may reduce the release of these cytokines in endometrial tissues. Therefore, PRP can be remedial in endometriosis. In different studies, however, IL1-B has been shown to increase, and similarly, also decrease with different L-PRPs and P-PRPs^52,53^. In our study, although the foci got smaller with P-PRP and L-PRP, we cannot say that PRP has a therapeutic effect on endometriosis since this reduction was also seen in the control group, and the difference in the reduction of volume was not significant. As Wang et al. have pointed out, the number of platelets, cytokines and factors in PRP may vary depending on how the PRP has been prepared, and these changes may explain the different outcomes in the literature^53^.

In our study, PRP derivatives were applied on the implant once in the form of an injection. Perhaps the application of the injections into the implant or intravenously, or simultaneously application of them with repeated doses at certain intervals could result in different and effective results. We can think of a limitation of our study that we did not histopathologically examine the similar implants in the first month in which the foci were found to have been formed. They have not been examined in many studies, either. We also performed an endometrial implantation, as in most past studies^22,23^. To the best of our knowledge, our current study is the first study of the relationship between PRP and endometriosis, which can be considered as a preliminary study. Although our results were not significant, it was promising that the PRP foci did not grow, and they did not stay the same — they shrank. Therefore, in order to investigate the effect of PRP, which has many important features, on endometriosis, there is a need for larger research studies which have different applications with different doses.

## Conclusion

In conclusion, the endometriosis foci were shrinking over time. This reduction was observed in all groups and was significant. However, the shrinking of endometriosis foci did not show any statistically significant difference among the groups. Moreover, there was no difference between the groups in terms of epithelial score, hemosiderin deposits, VEGF and total score. In other words, although both L-PRP and P-PRP generated more reduction in the endometriosis foci, they did not create any statistical differences.

## Acknowledgments

All authors have made significant contributions to the study.

## References

1. Kohl Schwartz AS, Wölfler MM, Mitter V, Rauchfuss M, Haeberlin F, Eberhard M. Et al. Endometriosis, especially mild disease: a risk factor for miscarriages. Fertil Steril. 2017; 108(5): 806–814.e2.

2. Marinho MCP, Magalhaes TF, Fernandes LFC, Augusto KL, Brilhante AVM, Bezerra LRPS. Quality of Life in Women with Endometriosis: An Integrative Review. J Womens Health (Larchmt). 2018; 27(3): 399–408.

3. Tremellen K, Thalluri V. Influence of Endometriosis on Assisted Reproductive Technology Outcomes: A Systematic Review and Meta-analysis. Obstet Gynecol. 2015; 125(6): 1498–9..

4. Rush G, Misajon R. Examining Subjective Wellbeing and Health-related quality of life in women with Endometriosis. Health Care Women Int. 39(3): 303–321.

5. Gordts S, Koninckx P, Brosens I. Pathogenesis of deep endometriosis. Fertil Steril. 2017; 108(6): 872–885.

6. Daraï E, Ploteau S, Ballester M, Bendifallah S. Pathogenesis, genetics and diagnosis of endometriosis. Presse Med. 2017; 46(12 Pt 1): 1156–1165.

7. Harada T, Iwabe T, Terakawa N. Role of cytokines in endometriosis. Fertil Steril. 2001; 76: 1–10.

8. McLaren J, Prentice A, Charnock-Jones DS, Smith SK. Vascular endothelial growth factor (VEGF) concentrations are elevated in peritoneal fluid of women with endometriosis. Hum Reprod. 1996; 11: 220–3.

9. Oosterlynck DJ, Meuleman C, Sobis H, Vandeputte M, Koninckx PR. Angiogenic activity of peritoneal fluid from women with endometriosis. Fertil Steril. 1993; 59: 778–82.

10. Tekin YB, Guven S, Kirbas A, Kalkan Y, Tumkaya L, Guven ESG. Is resveratrol a potential substitute for leuprolide acetate in experimental endometriosis? European Journal of Obstetrics & Gynecology and Reproductive Biology. 2015; 184: 1–6.

11. Uzunlar Ö, Ozyer Ş, Ustun YE, Moraloglu Ö, Gulerman HC, Caydere M. Et al. Effects of repeated propranolol administration in a rat model of surgically induced endometriosis European Journal of Obstetrics & Gynecology and Reproductive Biology. 2014; 182: 167–171.

12. Yılmaz S, Kaya E, Comakli S.Vitamin E (α tocopherol) attenuates toxicity and oxidative stress induced by aflatoxin in rats. Adv Clin Exp Med. 2017; 26(6): 907–917.

13. Pala Ş, Atilgan R, Kuloğlu T, Kara M, Başpinar M, Can B, et al. Protective effects of vitamin C and vitamin E against hysterosalpingography-induced epithelial degeneration and proliferation in rat endometrium. Drug Des Devel Ther. 2016; 15(10): 4079–4089.

14. Molavi M, Razi M, Cheraghi H, Khorramjouy M, Ostadi A, Gholirad S. Protective effect of vitamin E on cypermethrin-induced follicular atresia in rat ovary: Evidence for energy dependent mechanism. Vet Res Forum. 2016; 7(2): 125–132.

15. Hoeferlin LA, Huynh QK, Mietla JA, Sell SA, Tucker J, Chalfant CE, et al. The lipid portion of activated platelet rich plasma significantly contributes to its wound healing properties. Adv Wound Care. 2015; 4(2): 100–109.

16. Demirbag S, Cetinkursun S, Tasdemir U, Ozturk H, Pekcan M, Yesildaglar N. Comparison of hyaluronate/carboxymethylcellulose membrane and melatonin for prevention of adhesion formation in a rat model. Hum Reprod. 2005; 20(7): 2021–2024.

17. Ferrando J, García-García SC, González-de-Cossío AC, Bou L, Navarra E. A Proposal of an Effective Platelet-rich Plasma Protocol for the Treatment of Androgenetic Alopecia. Int J Trichology. 2017; 9(4):165–170..

18. Martini LI, Via AG, Fossati C, Randelli F, Randelli P, Cucchi D. Single Platelet-Rich Plasma Injection for Early Stage of Osteoarthritis of the Knee. Joints. 2017; 5(1):2–6.

19. Unlu MC, Kivrak A, Kayaalp ME, Birsel O, Akgun I. Peritendinous injection of platelet-rich plasma to treat tendinopathy: A retrospective review. Acta Orthop Traumatol Turc. 2017; 51(6): 482–487.

20. Yang WY, Han YH, Cao XW, Pan JK, Zeng LF, Lin JT, et al. Platelet-rich plasma as a treatment for plantar fasciitis: A meta-analysis of randomized controlled trials. Medicine (Baltimore). 2017; 96(44): e8475.

21. Jia J, Wang S, Ma L, Yu J, Guo Y, Wang C. The Differential Effects of Leukocyte-Containing and Pure Platelet-Rich Plasma on Nucleus Pulposus-Derived Mesenchymal Stem Cells: Implications for the Clinical Treatment of Intervertebral Disc Degeneration. Stem Cells Int. 2018; 2018: 7162084.

22. Vernon MW, Wilson EA. Studies on the surgical induction of endometriosis in the rat. Fertil Steril. 1985; 44:684–694

23. Uygur D, Aytan H, Zergeroglu S, Batioglu S. Leflunomide—an immunomodulator— induces regression of endometrial explants in a rat model of endometriosis. J Soc Gynecol Investig. 2006; 13: 378–383

24. Islimye M, Kilic S, Zulfikaroglu E, Topcu O, Zergeroglu S, Batioglu S. Regression of endometrial autografts in a rat model of endometriosis treated with etanercept. European Journal of Obstetrics & Gynecology and Reproductive Biology. 2011; 159: 184–189.

25. Oz M, Cetinkaya N, Bas S, Korkmaz E, Ozgu E, Terzioglu GS. et al. A randomized controlled experimental study of the efficacy of platelet-rich plasma and hyaluronic acid for the prevention of adhesion formation in a rat uterine horn model. Arch Gynecol Obstet. 2016; 294: 533–540.

26. Xu Z, Yin W, Zhang Y, Qi X, Chen Y, Xie X, et al. Comparative evaluation of leukocyte- and platelet-rich plasma and pure platelet-rich plasma for cartilage regeneration. Sci Rep. 2017; 7: 43301.

27. Ata B, Uncu G. Impact of endometriomas and their removal on ovarian reserve. Curr Opin Obstet Gynecol. 2015; 27: 235–241.

28. Somigliana E, Berlanda N, Benaglia L, Vigano P, Vercellini P, Fedele L. Surgical excision of endometriomas and ovarian reserve: a systematic review on serum antimullerian hormone level modifications. Fertil Steril. 2012; 98: 1531–1538

29. Kondo W, Bourdel N, Tamburro S, Cavoli D, Jardon K, Rabischong B. et al Complications after surgery for deeply infiltrating pelvic endometriosis. BJOG. 2011; 118(3): 292–8.

30. Zwiep T, Humphrey R, Fortin D, Inculet RI, Malthaner RA. Autologous platelet rich plasma and concentrated platelet poor plasma are safe in patients requiring lobectomies but do not reduce the duration of air leak: a randomized controlled trial. Ann SurgInt. 2016; 2(2): ASI-2-011.

31. Le ADK et al. Platelet-Rich Plasma. Clin Sports Med. 2019; 38(1): 17–44.

32. Yildirim G, Attar R, Ficicioglu C, Karateke A, Ozkan F, Yesildaglar N. Etanercept causes regression of endometriotic implants in a rat model. Arch Gynecol Obstet. (2011) 283: 1297–1302

33. Bedaiwy S. Gupta J. Noriega J. Brainard A. Agarwal T. Falcone Diagnostic value of hemosiderin laden macrophages in histologically proven endometriosis. Fertility and Sterility. 2008 Sep; 90: Supplement page S438

34. Bedaiwy MA, Noriega J, Abdel Aleem M, Gupta S, AbulHassan AM, Brainard J. et al. Evaluation of peritoneal fluid hemosiderin-laden macrophages in biopsy-proven endometriosis. Anal Quant Cytol Histol. 2012; 34(1): 23–7.

35. Zadehmodarres S, Salehpour S, Saharkhiz N, Nazari L. Treatment of thin endometrium with autologous platelet-rich plasma: a pilot study. JBRA Assisted Reproduction. 2017; 21(1): 54–56

36. Chang Y, Li J, Chen Y, Wei L, Yang X, Shi Y, Liang X. Autologous platelet-rich plasma promotes endometrial growth and improves pregnancy outcome during in vitro fertilization. Int J Clin Exp Med. 2015; 8(1): 1286–1290.

37. Farimani M, Poorolajal J, Rabiee S, Bahmanzadeh M. Successful pregnancy and live birth after intrauterine administration of autologous platelet-rich plasma in a woman with recurrent implantation failure: A case report. Int J Reprod BioMed. 2017; 15 (12): 803–806.

38. Farimani M, Bahmanzadeh M, Poorolajal J. A New Approach Using Autologous Platelet-Rich Plasma to Treat Infertility and To Improve Population Replacement Rate. J Res Health Sci. 2016; 16(3): 172–173.

39. Reghini MFS, Ramires Neto C, Segabinazzi LG, Castro Chaves, Dell’Aqua Cd, Bussiere MCC.et al. Inflammatory Response in Chronic Degenerative Endometritis Mares Treated with Platelet-Rich Plasm. Theriogenology. 2016; 15; 86(2): 516–22.

40. Anitua E, Andia I, Ardanza B, Nurden P, Nurden AT. Autologous platelets as a source of proteins for healing and tissue regeneration. Thromb Haemost. 2004; 91: 4–15.

41. Gentile P, Orlandi A, Scioli MG, Di Pasquali C, Bocchini I, Cervelli V. Concisereview: adipose-derived stromal vascular fraction cells and platelet-richplasma: basic and clinical implications for tissue engineering therapies inregenerative surgery. Stem Cells Transl Med. 2012; 1: 230–6.

42. Başbuğ M, Arıkanoğlu Z. Formation And Clinical İmportance Of Postoperative Peritoneal Adhesions. J Clin Exp Invest. 2010; 1(2): 134–137

43. Zhang XL, Shi KQ, Jia PT, Jiang LH, Liu YH, Chen X.et al. Effects of platelet-rich plasma on angiogenesis and osteogenesis-associated factors in rabbits with avascular necrosis of the femoral head. Eur Rev Med Pharmacol Sci. 2018; 22(7): 2143–2152.

44. Bastu E, Mutlu MF, Yasa C, Attar NE. Endometriosis and immunology. 2013; 1(2): 54–62

45. Kupker W, Schultze-Mosgau A, Diedrich K. Paracrine changes in the peritoneal environment of women with endometriosis. Hum Reprod Update. 2000; 4(5): 719–723.

46. Ergenoğlu AM, Yeniel AÖ, Erbaş O, Aktuğ H, Yildirim N, Ulukuş M. et al. Regression of endometrial implants by resveratrol in an experimentally induced endometriosis model in rats. Reprod Sci. 2013; 20(10): 1230–6.

47. Imai S, Kumagai K, Yamaguchi Y, Miyatake K, Saito T. Platelet-Rich Plasma Promotes Migration, Proliferation, and the Gene Expression of Scleraxis and Vascular Endothelial Growth Factor in Paratenon-Derived Cells In Vitro. Sports Health. 2012; 40(5): 1035–45

48. Canbeyli İD, Akgun RC, Sahin O, Terzi A, Tuncay İC. Platelet-rich plasma decreases fibroblastic activity and woven bone formation with no significant immunohistochemical effect on long-bone healing: An experimental animal study with radiological outcomes. J Orthop Surg (Hong Kong). 2018; 26(3): 2309499018802491.

49. Rodrigues BL, Montalvão SA, Cancela RB, Silva FA, Urban A, Huber SC. et al. “Treatment of male pattern alopecia with platelet-rich plasma: a double blind controlled study with analysis of platelet number and growth factor levels”. Journal of the American Academy of Dermatology 2019; 80(3): 694–700.

50. Chen CH, Cao Y, Wu YF, Bais AJ, Gao JS, Tang JB. Tendon healing in vivo: gene expression and production of multiple growth factors in early tendon healing period. J Hand Surg. 2008; 33: 1834–42.

51. Marini GM, Perrini C, Esposti P, Corradetti B, Bizzaro D, Riccaboni P, et. al. Effects of platelet-rich plasma in a model of bovine endometrial inflammation in vitro Maria Giovanna. Reproductive Biology and Endocrinology. 2016; 14(1): 58.

52. Menchisheva Y, Mirzakulova U, Yui R. Use of platelet-rich plasma to facilitate wound healing. Int Wound J. 2019; 16(2): 343–353.

53. Wang K, Li Z, Li J, Liao W, Qin Y, Zhang N.et al.Optimization of the Platelet-Rich Plasma Concentration for Mesenchymal Stem Cell Applications. Tissue Eng Part A. 2019; 25(5-6): 333–351.

